## Supplemental Figures for "The genomic landscape of transposable elements in yeast hybrids is shaped by structural variation and genotype-specific modulation of transposition rate"

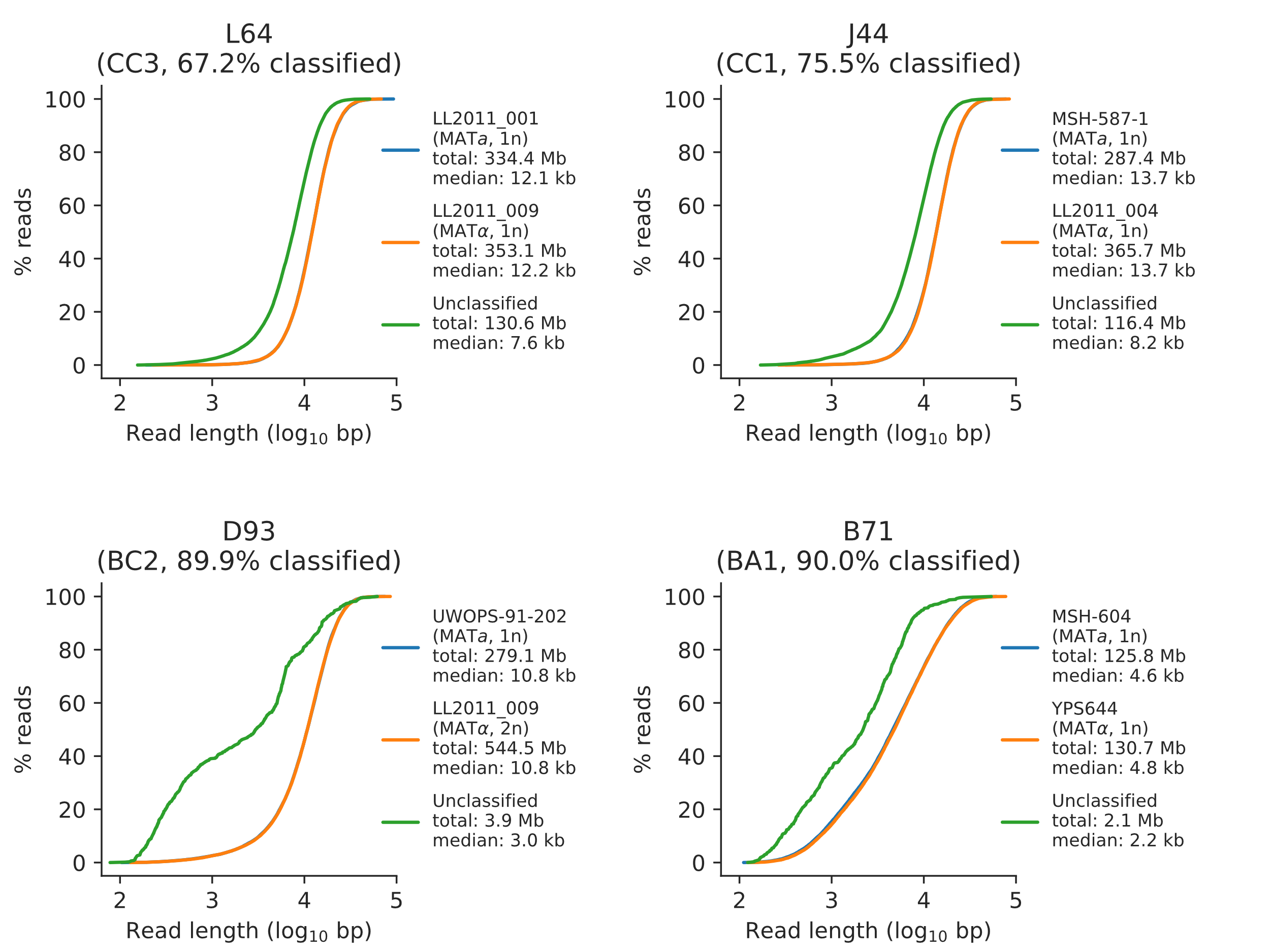

### Supplemental Figure S1. Read length distributions of classified and unclassified reads for a representative sample of four MA lines long-read libraries.

Cumulative length distributions for reads classified as the *MAT*a parental subgenome, reads classified as the *MAT*α parental subgenome, and unclassified reads are shown in blue, orange and green respectively. Subgenome ploidy (if applicable), total bases and median read length are shown for each category.

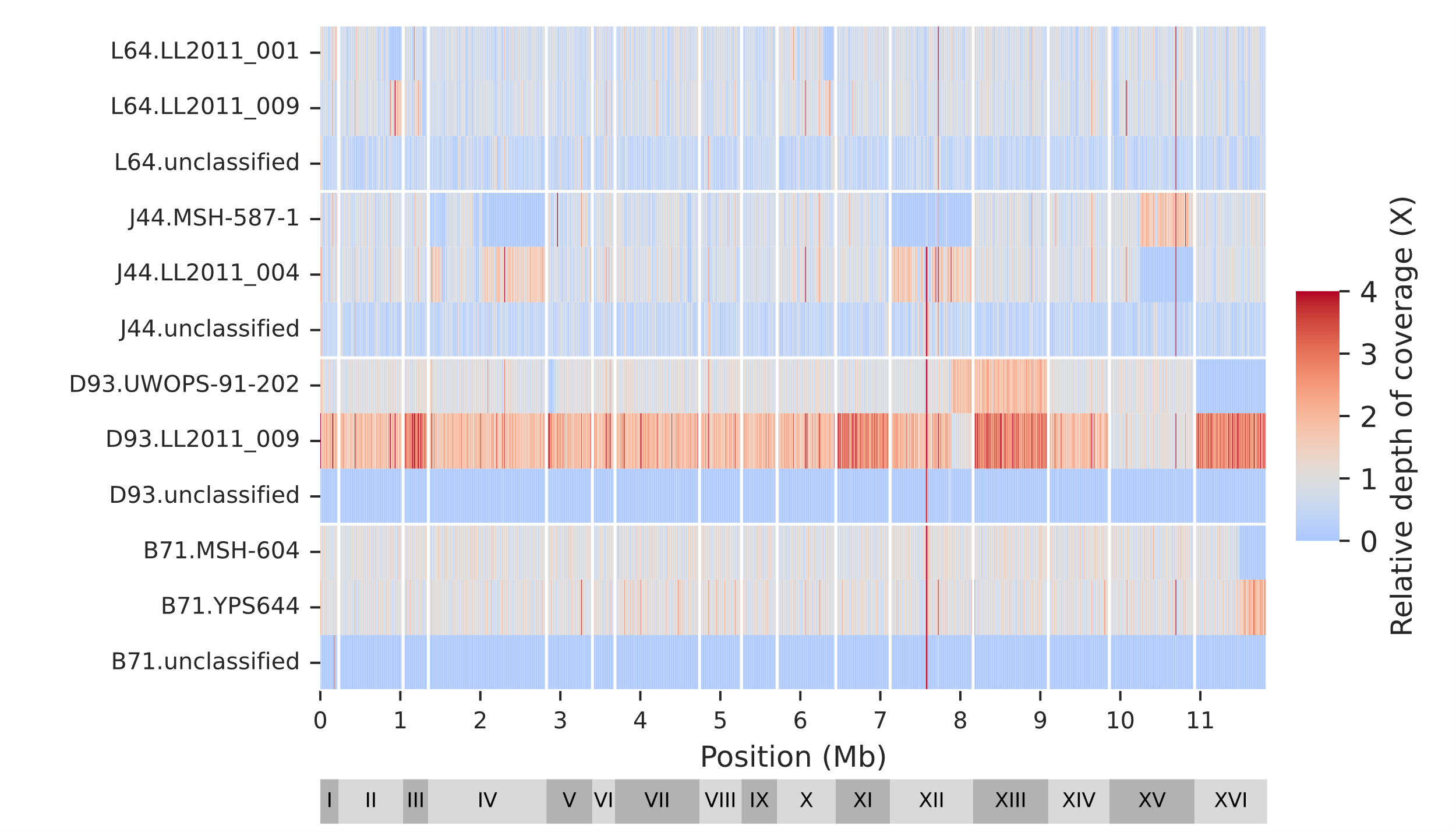

### Supplemental Figure S2. Genomic distribution of classified and unclassified reads.

The MA lines long-read libraries are the same as shown in Supplemental Figure S1.

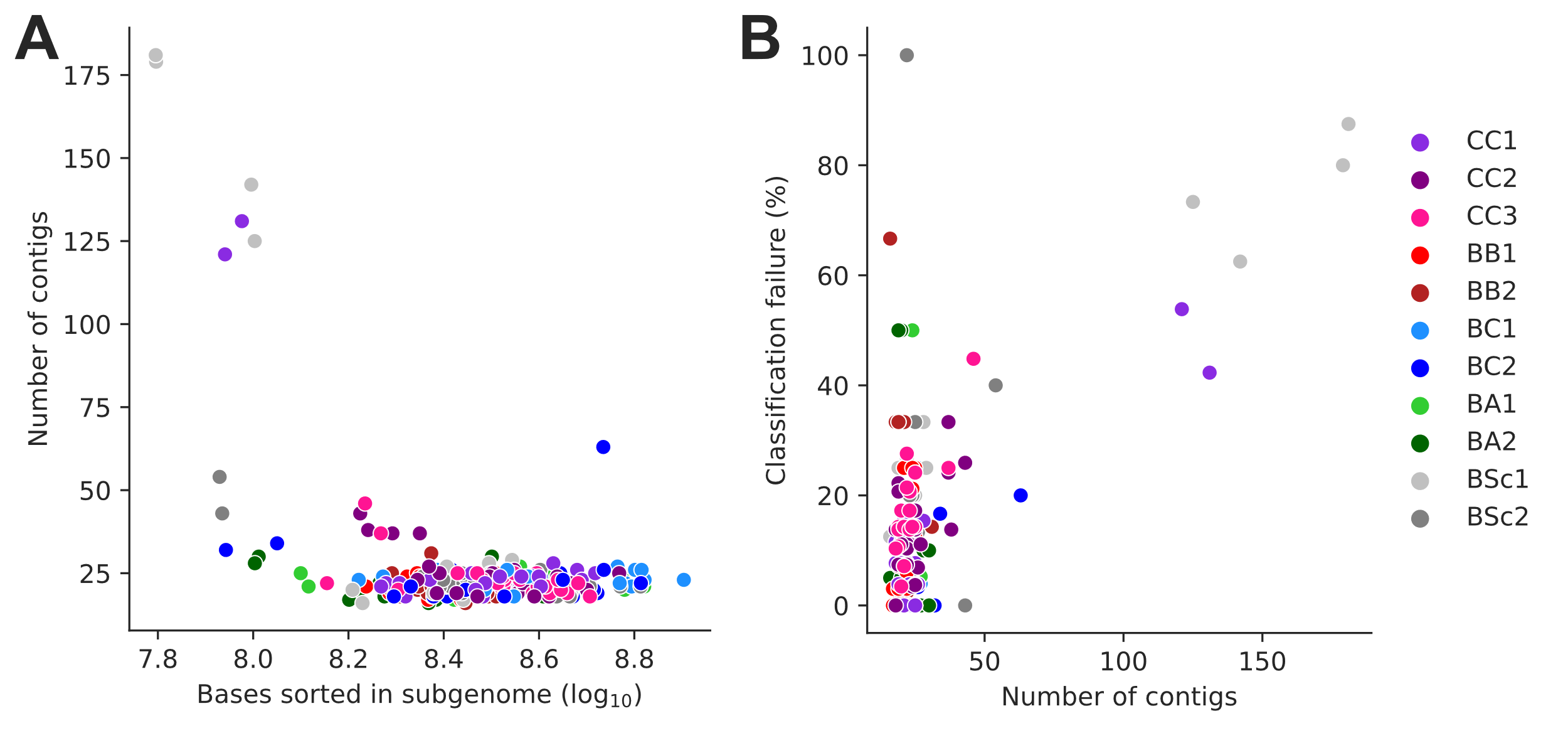

### Supplemental Figure S3. Subgenome-level assemblies of low contiguity.

**(A)** Assembly contiguity (number of contigs) is largely determined by the size of the corresponding sorted long-read library. **(B)** Assembly contiguity largely determines Ty loci classification failure rate.

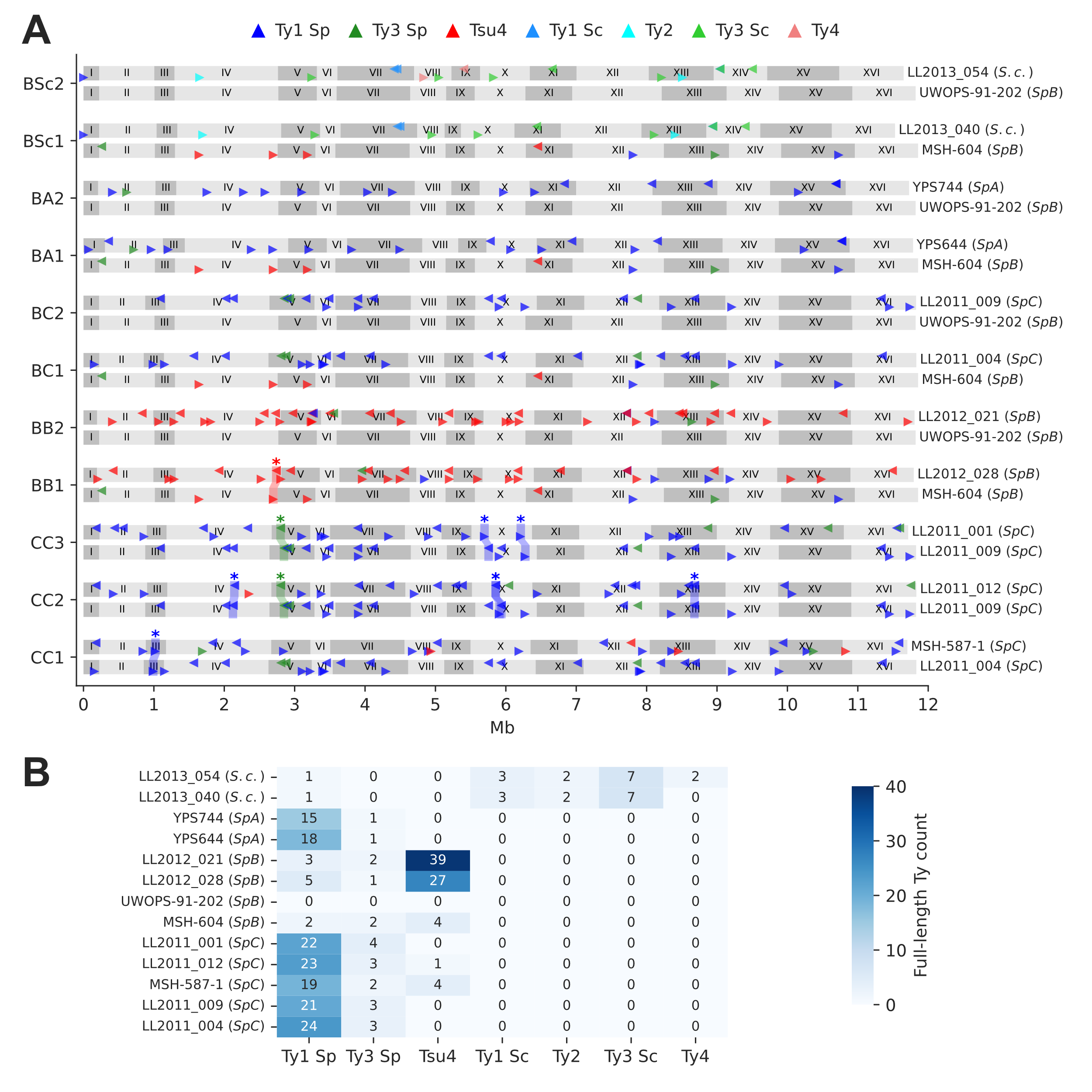

### Supplemental Figure S4. Ty element annotations in the parental genomes.

**(A)** Genome maps show the annotations of full-length Ty elements for the two parental genomes of each cross. Triangle orientation indicates whether the annotation is on the plus (right pointing) or minus (left pointing) chromosome strand. Shaded areas labelled with stars highlight pairs of parental annotations that were inferred to be orthologous based on whole-genome alignments. Some orthologous pairs are also linked to inversions or duplications. **(B)** Count of full-length annotations for each genome.

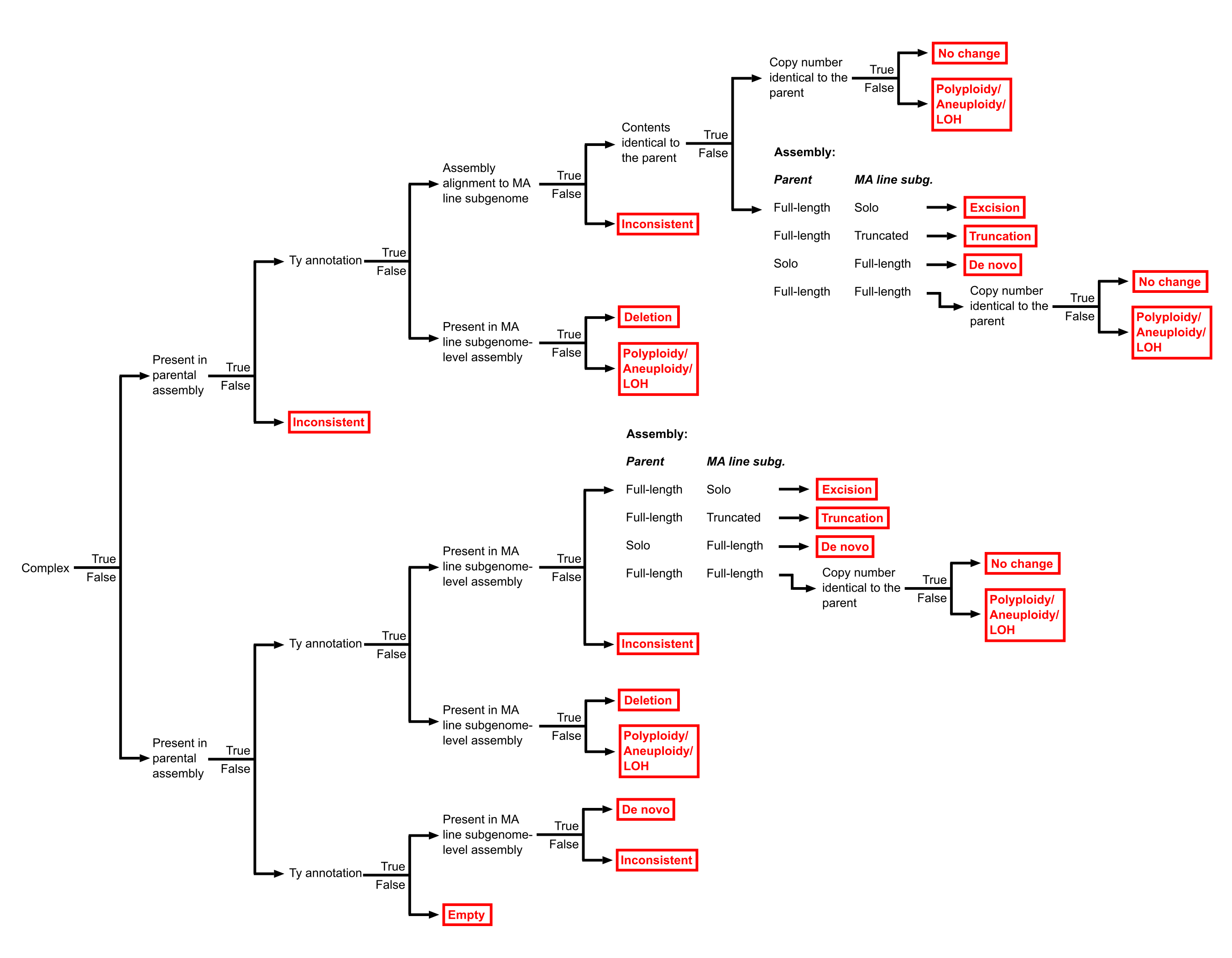

### Supplemental Figure S5. Hierarchy of binary rules used for the classification of Ty loci into SV types.

The classification outcomes, including the different SV types, are shown in red.

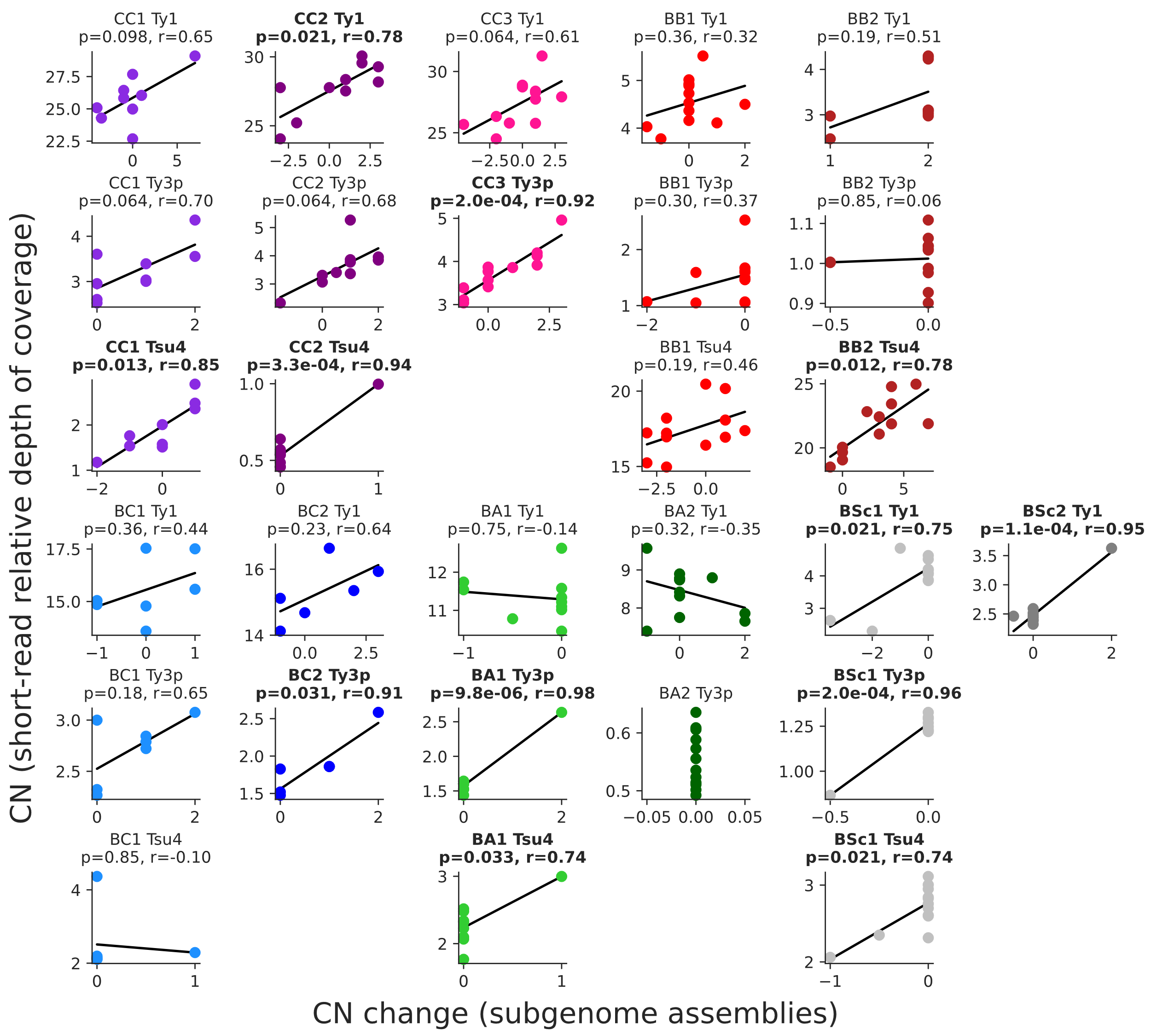

### Supplemental Figure S6. Correlations between the estimations of full-length Ty copy numbers in the MA lines genomes.

Copy number changes from the annotation of subgenome assemblies (this study) are shown on the X-axis, while copy numbers estimated from relative depth of coverage of short-read libraries after evolution by MA (Hénault et al., 2020) are shown on the Y-axis. Where applicable, FDR-corrected P-values and Pearson correlation coefficients are shown, with significant correlation in bold.

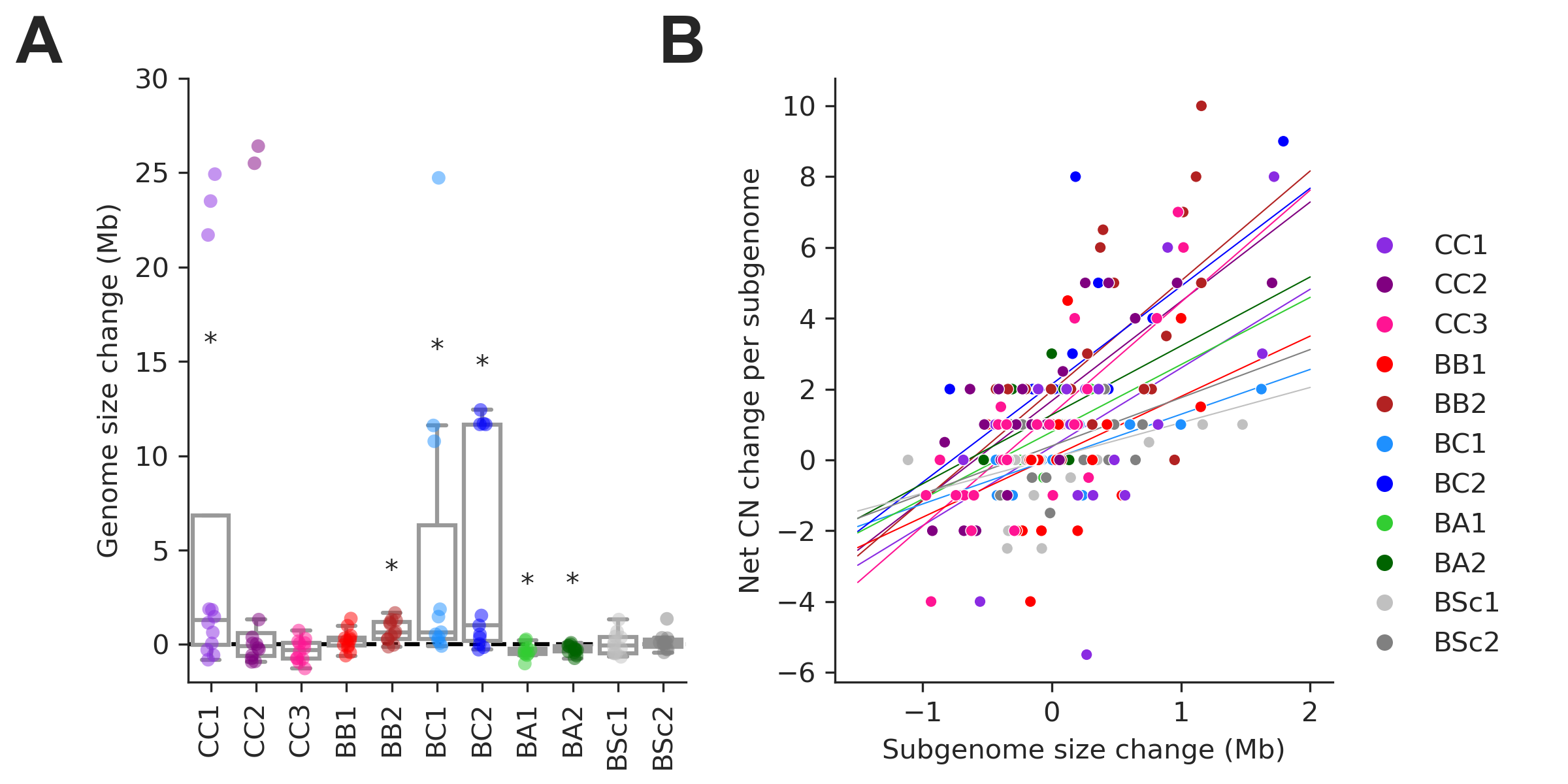

### Supplemental Figure S7. Changes in Ty CN are correlated with genome expansion.

**(A)** Distributions of genome size change for individual MA lines. Symbols indicate FDR-corrected P-values for Wilcoxon signed-rank tests for deviations from zero (*: p≤0.05, **: p≤0.01). **(B)** Correlations between net Ty CN change per MA line subgenome and subgenome size change. Background lines represent the fit of a linear mixed effect model including random intercepts and slopes for each MA cross.

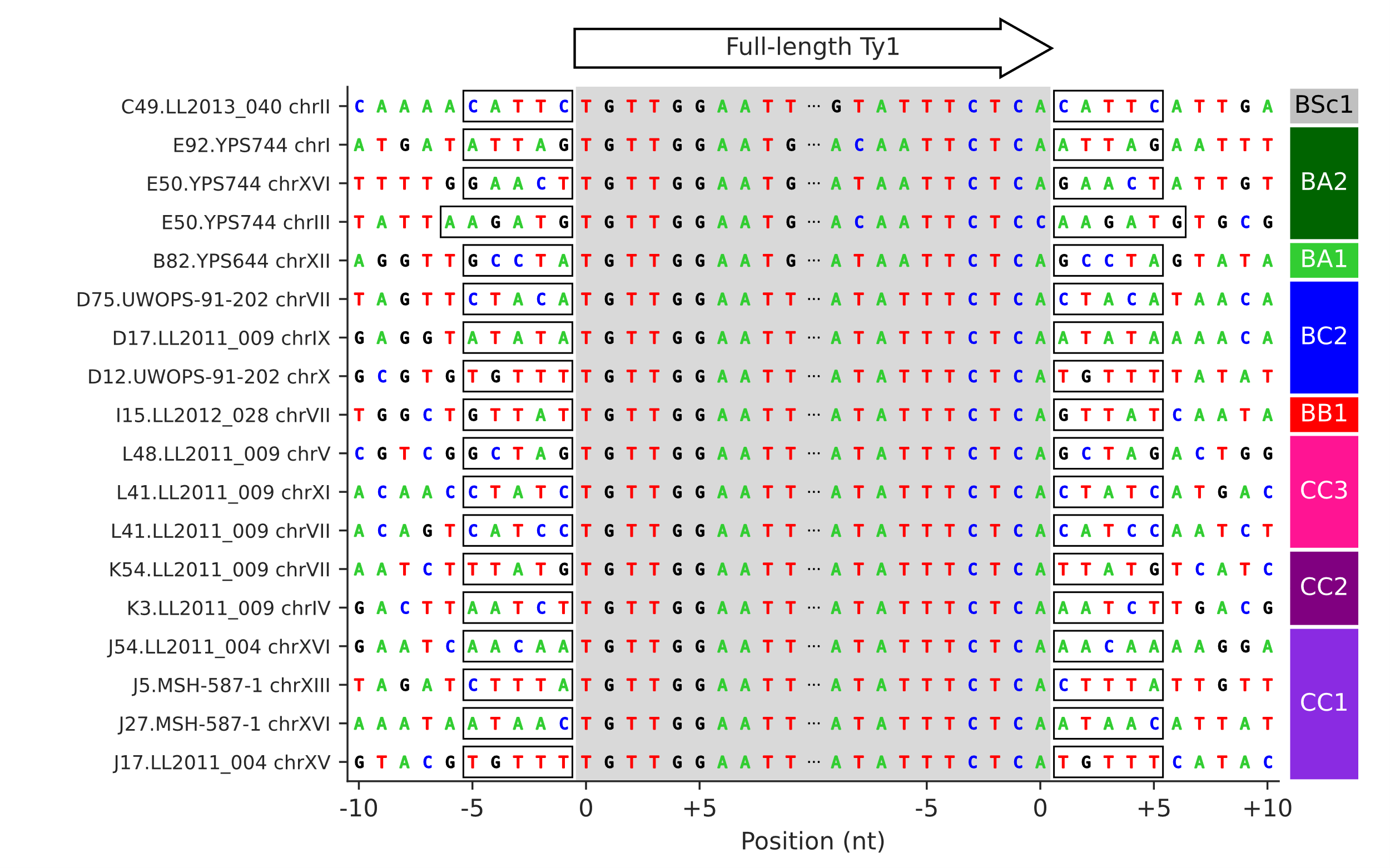

### Supplemental Figure S8. Target site duplication (TSD) sequences of the 18 confident Ty1 de novo retrotransposition events.

The grey shaded area corresponds to the 5’ to 3’ sequence of full-length Ty1 elements. Flanking sequences correspond to the 10 nucleotides immediately upstream and downstream of the insertion site. Identical 5’ and 3’ TSDs are highlighted by frames.

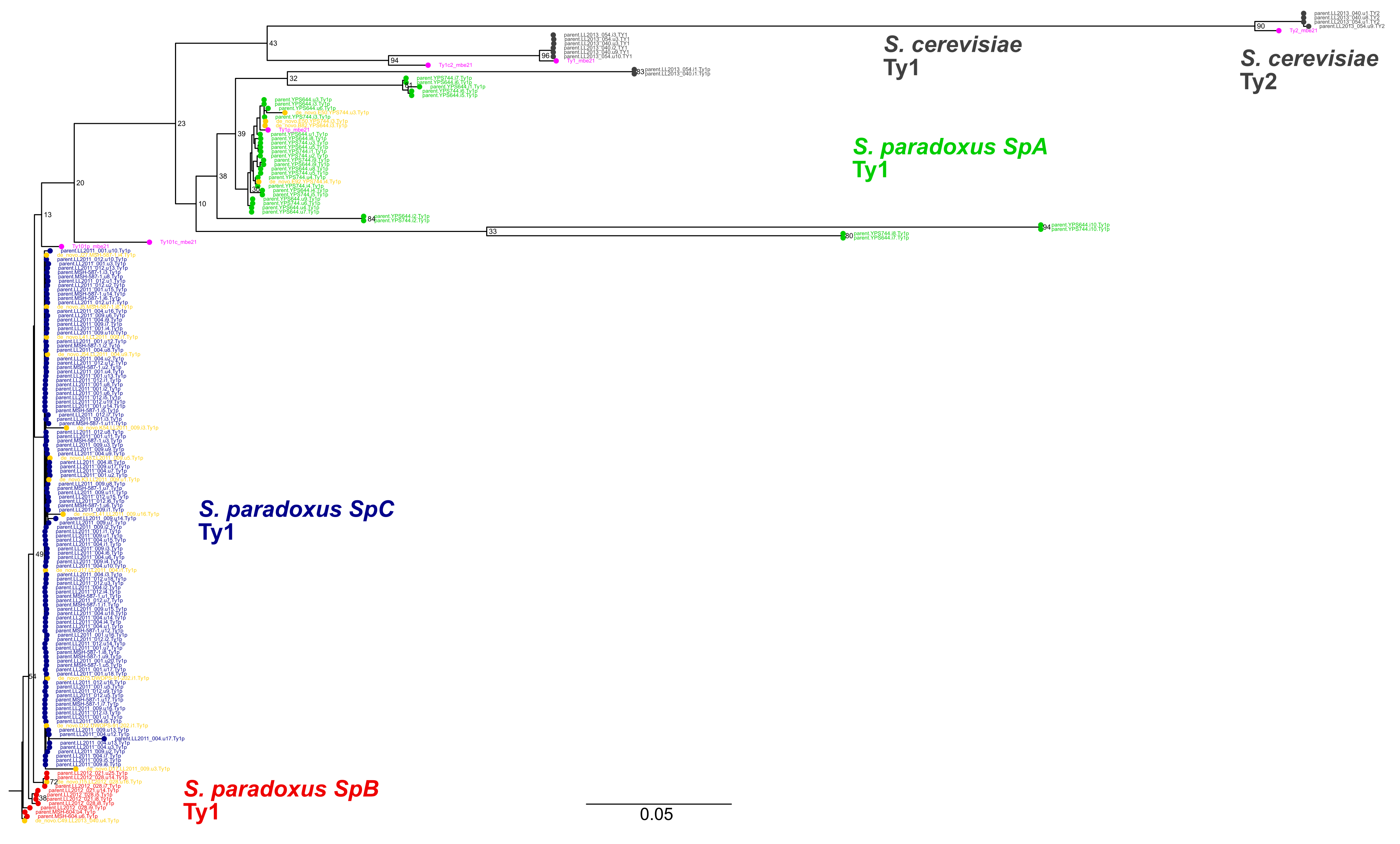

### Supplemental Figure S9. Maximum likelihood phylogenetic tree comprising the full-length parental and de novo Ty1 and Ty2 elements.

De novo retrotransposition events are shown in yellow. Representative copies of the Ty1 and Ty2 subfamilies defined by Bleykasten-Grosshans et al. (2021) are shown in fuchsia. Bootstrap support values are shown as percentages for the selected major branches. The scale is in substitutions per site.

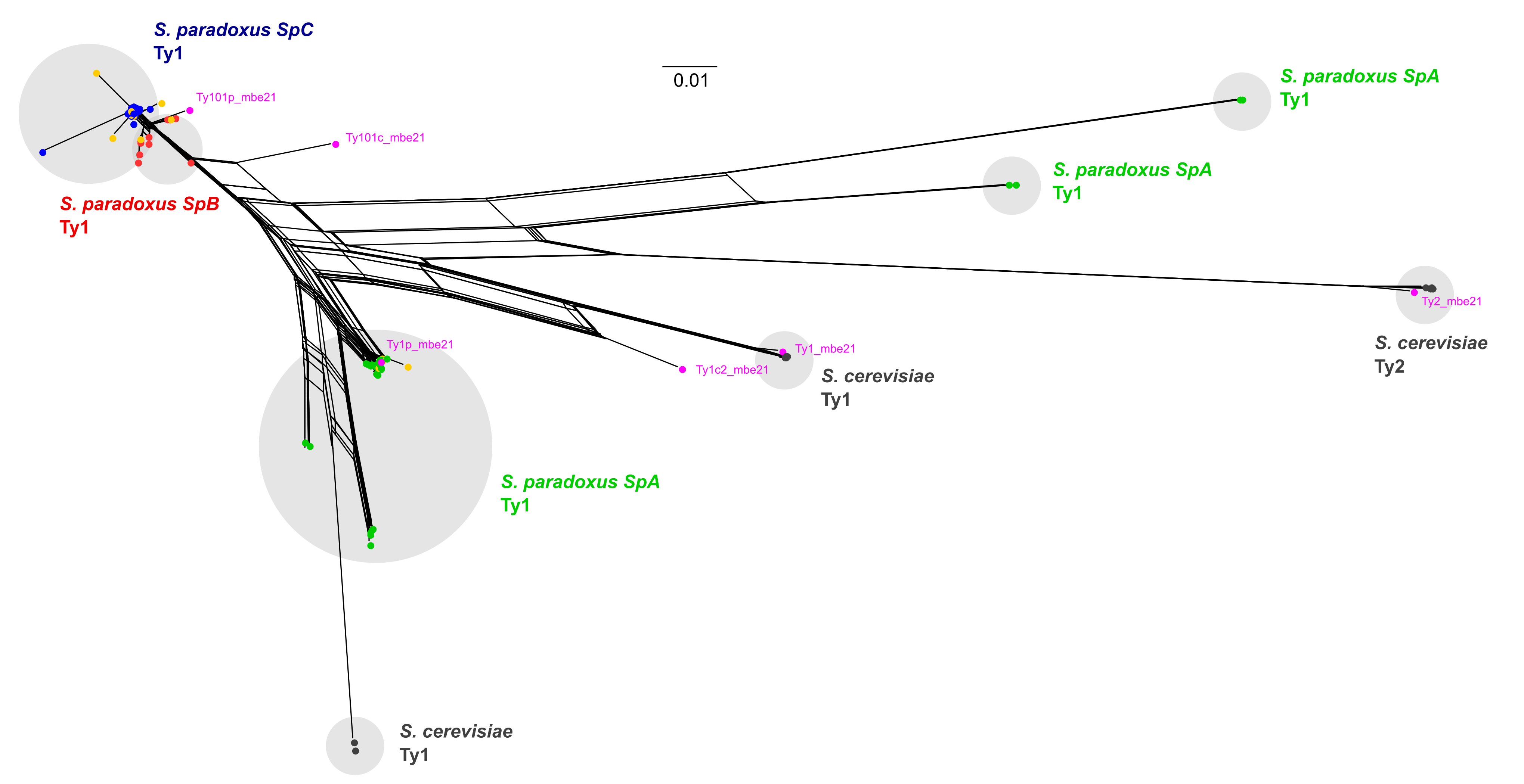

### Supplemental Figure S10. Phylogenetic network comprising the full-length parental and de novo Ty1 and Ty2 elements.

De novo retrotransposition events are shown in yellow. Representative copies of the Ty1 and Ty2 subfamilies defined by Bleykasten-Grosshans et al. (2021) are shown in fuchsia. The scale is in substitutions per site.

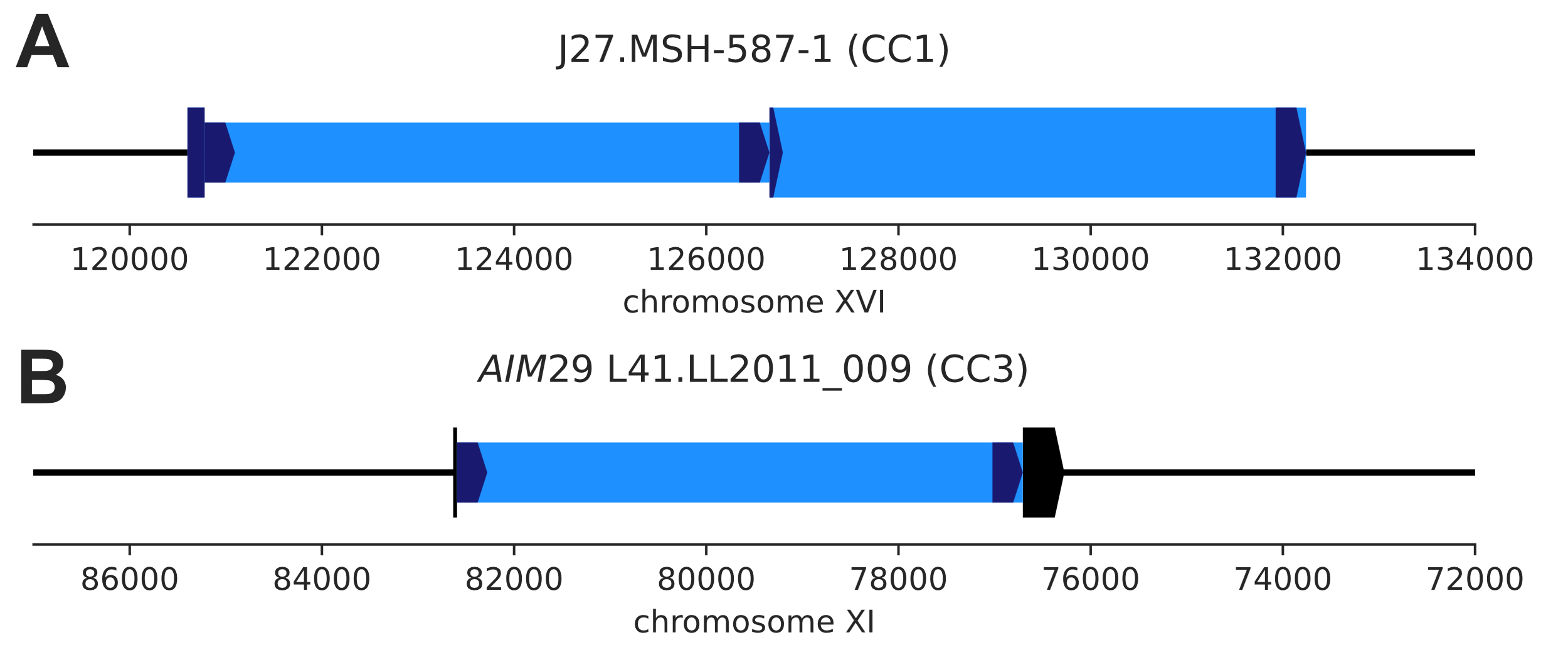

### Supplemental Figure S11. De novo Ty1 transposition events disrupting parental genomic features.

**(A)** De novo Ty1 insertion (narrow box) into an existing full-length Ty1 element (wide box) in the MSH-587-1 (*SpC*) subgenome of MA line J27 (CC1). **(B)** De novo Ty1 insertion (narrow box) into the host gene *AIM29* (YKR074W, wide box) in the LL2011_009 (*SpC*) subgenome of MA line L41 (CC3).

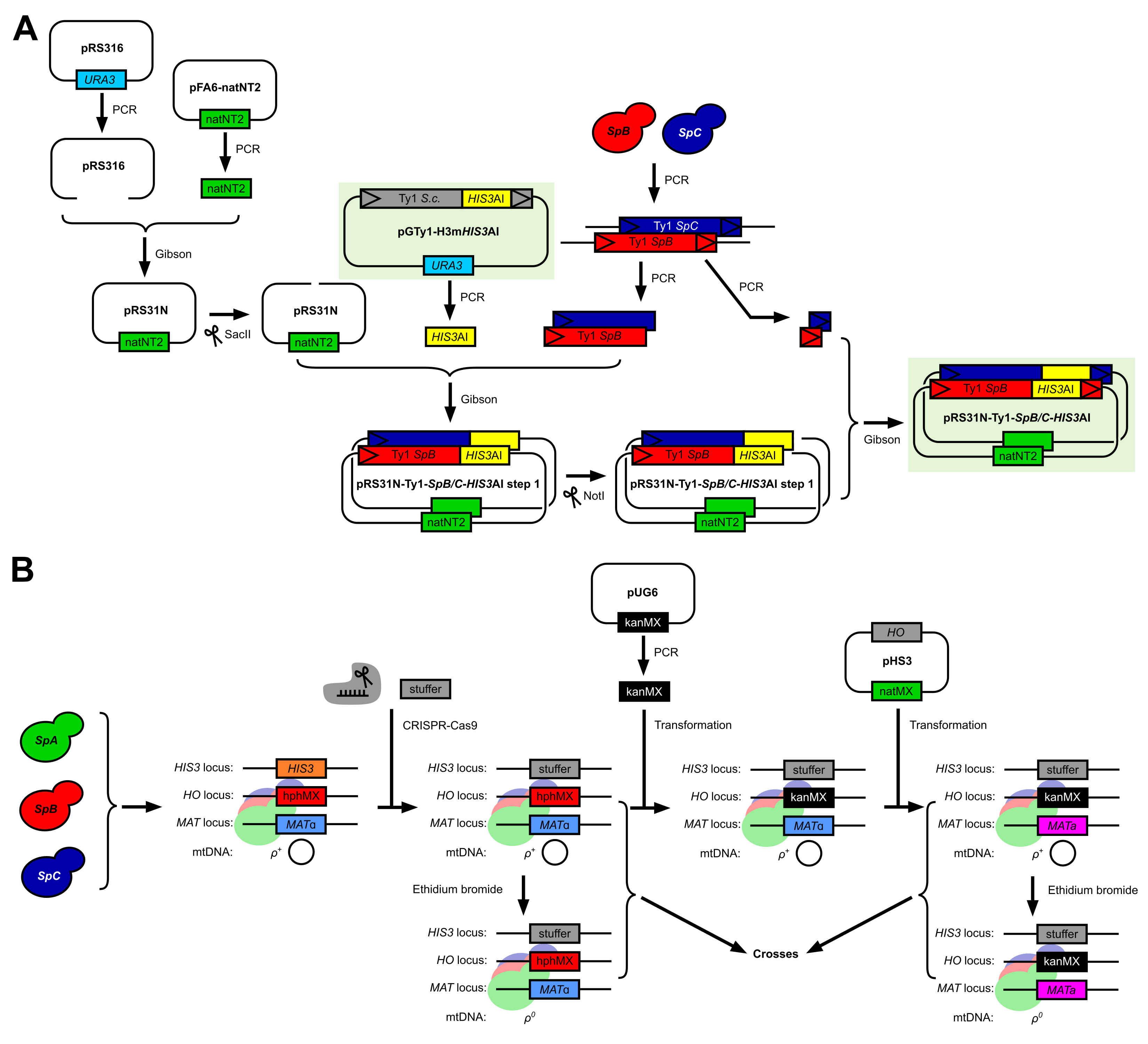

### Supplemental Figure S12. Methodology for the adaptation of the in vivo *S. cerevisiae* Ty1 retrotransposition assay to *S. paradoxus* Ty1 variants and genetic backgrounds.

**(A)** Construction of the pRS316-Ty1-*SpB*-*HIS3*AI and pRS316-Ty1-*SpC*-*HIS3*AI plasmids with *SpB* and *SpC* Ty1 sequence variants, respectively. **(B)** Construction of a panel of histidine-auxotroph *S. paradoxus* haploid strains for the generation of combinatorial crosses with reciprocal mtDNA inheritance.

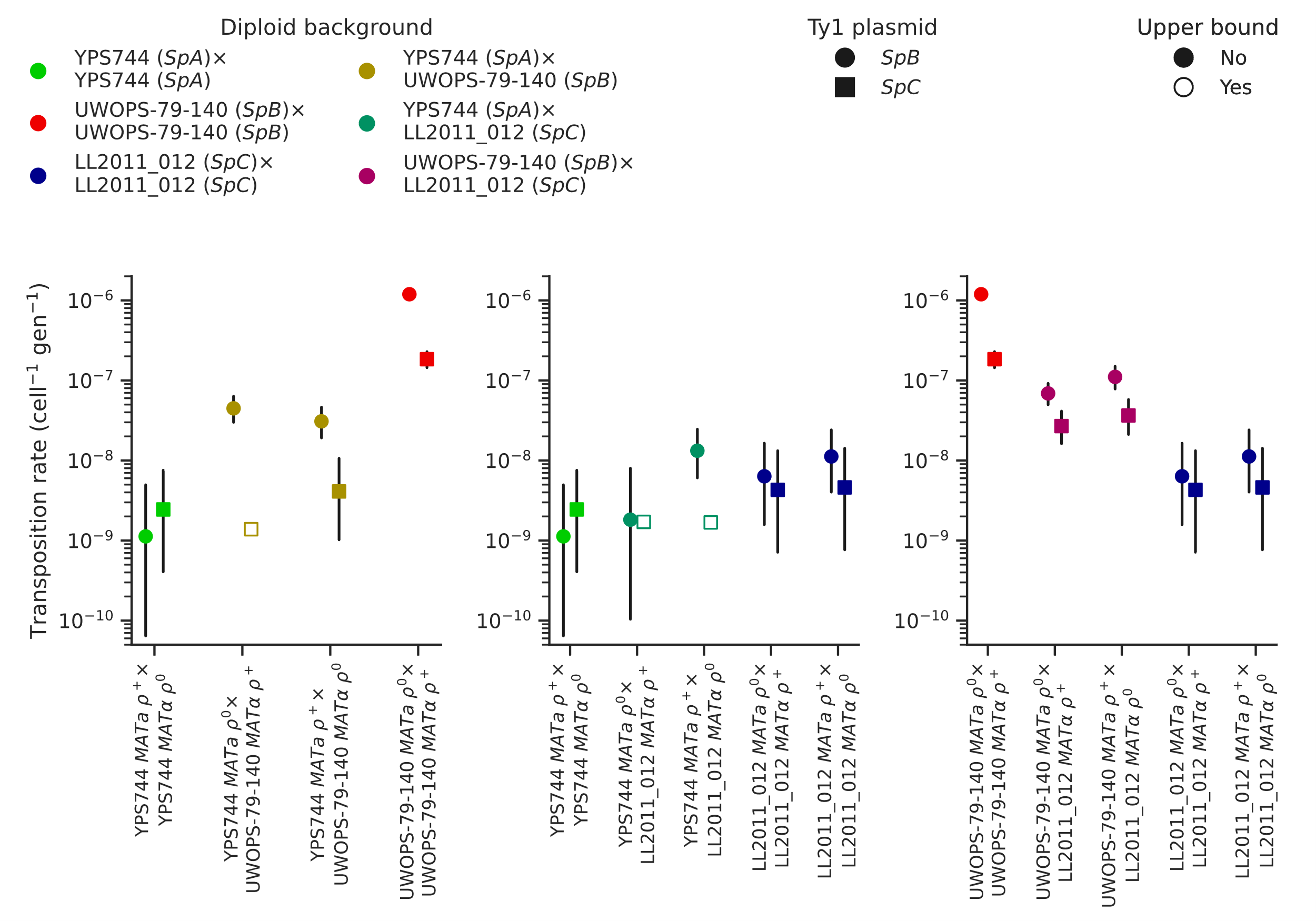

### Supplemental Figure S13. Ty1 transposition rates in diploid homozygous backgrounds of natural populations of *S. paradoxus* and their hybrids.

Transposition rates were measured in three diploid *S. paradoxus* backgrounds and their hybrids. Black bars indicate 95% confidence intervals. Empty symbols represent upper bound transposition rate estimates obtained by artificially inflating the mutant count by one when zero was observed. Marker shapes indicate the sequence variant of the tester Ty1 element, either *SpB* (circles) or *SpC* (squares).
